# BMN 250, a fusion of lysosomal alpha-N-acetylglucosaminidase with IGF2, exhibits different patterns of cellular uptake into critical cell types of Sanfilippo syndrome B disease pathogenesis

**DOI:** 10.1101/466078

**Authors:** Gouri Yogalingam, Amanda R Luu, Heather Prill, Melanie J. Lo, Bryan Yip, John Holtzinger, Terri Christianson, Mika Aoyagi-Scharber, Roger Lawrence, Brett E. Crawford, Jonathan H. LeBowitz

## Abstract

Sanfilippo syndrome type B (Sanfilippo B; Mucopolysaccharidosis type IIIB) occurs due to genetic deficiency of lysosomal alpha-N-acetylglucosaminidase (NAGLU) and subsequent lysosomal accumulation of heparan sulfate (HS), which coincides with devastating neurodegenerative disease. Because NAGLU expressed in Chinese hamster ovary cells is not mannose-6-phosphorylated, we developed an insulin-like growth factor 2 (IGF2)-tagged NAGLU molecule (BMN 250; tralesinidase alfa) that binds avidly to the IGF2 / cation-independent mannose 6-phosphate receptor (CI-MPR) for glycosylation independent lysosomal targeting. BMN 250 is currently being developed as an investigational enzyme replacement therapy for Sanfilippo B. Here we distinguish two cellular uptake mechanisms by which BMN 250 is targeted to lysosomes. In normal rodent-derived neurons and astrocytes, the majority of BMN250 uptake over 24 hours reaches saturation, which can be competitively inhibited with IGF2, suggestive of CI-MPR-mediated uptake. K_uptake_, defined as the concentration of enzyme at half-maximal uptake, is 5 nM and 3 nM in neurons and astrocytes, with a maximal uptake capacity (V_max_) corresponding to 764 nmol/hr/mg and 5380 nmol/hr/mg, respectively. Similar to neurons and astrocytes, BMN 250 uptake in Sanfilippo B patient fibroblasts is predominantly CI-MPR-mediated, resulting in augmentation of NAGLU activity with doses of enzyme that fall well below the K_uptake_ (5 nM), which are sufficient to prevent HS accumulation. In contrast, uptake of the untagged recombinant human NAGLU (rhNAGLU) enzyme in neurons, astrocytes and fibroblasts is negligible at the same doses tested. In microglia, receptor-independent uptake, defined as enzyme uptake resistant to competition with excess IGF2, results in appreciable lysosomal delivery of BMN 250 and rhNAGLU (V_max_=12,336 nmol/hr/mg and 5469 nmol/hr/mg, respectively). These results suggest that while receptor-independent mechanisms exist for lysosomal targeting of rhNAGLU in microglia, BMN 250, by its IGF2 tag moiety, confers increased CI-MPR-mediated lysosomal targeting to neurons and astrocytes, two additional critical cell types of Sanfilippo B disease pathogenesis.

## INTRODUCTION

Heparan sulfate (HS)–containing proteoglycans in the extracellular matrix and at the cell surface play important roles in the regulation of protease activity, growth factor signaling and cell surface receptor-mediated endocytosis of various ligands (Mulloy and Rider et al., 2006; Christianson and Belting, 2014). While little is understood regarding how these macromolecules contribute to disease one clue to their biological importance lies in a devastating group of inherited diseases involving impaired turnover of HS in lysosomes. Mucopolysaccharidosis III (MPS III; Sanfilippo syndrome) includes four biochemically distinct lysosomal storage diseases, each characterized by deficiency of a lysosomal enzyme involved in the step-wise degradation of HS (Neufeld and Muenzer, 2001). Sanfilippo B exhibits an autosomal recessive mode of inheritance and arises from mutations in the *NAGLU* gene, which encodes α-N-acetylglucosaminidase (NAGLU). In its severest form Sanfilippo B patients exhibit non-detectable or very low levels of NAGLU activity in their lysosomes, which coincides with lysosomal HS accumulation in the brain and ultimately leads to severe neurodegenerative disease and early death (Andrade et al., 2015; Yogalingam & Hopwood 2001). However, little is understood regarding the mechanism by which accumulation of HS in lysosomes leads to neurodegenerative disease in Sanfilippo syndrome. Studies in a mouse model of Sanfilippo B suggest that neurons and astrocytes of the Sanfilippo B brain may be compromised as a consequence of elevated levels of HS (Li et al., 2002, Ohmi et al., 2009). HS accumulation in microglia is also thought to contribute to neurodegeneration associated with Sanfilippo B (Ohmi et al., 2003; Archer et al., 2014). Developing therapeutic approaches to augment NAGLU activity in these critical cell types of neurodegenerative disease pathogenesis in Sanfilippo B may therefore be paramount for successful clearance of accumulated substrate, subsequently leading to correction of underlying pathology and stabilization of the disease.

In most cases of Sanfilippo B and other MPS disorders, the entire range of clinical phenotypes ranging from severe to relatively mild are clustered within a narrow range of residual lysosomal enzyme activity, from 0-20% of normal control levels (Yogalingam et al., 2000; Yogalingam et al., 1998; Crawley et al., 1998; Clark et al., 2018). These genotype-phenotype correlations suggest that reduced levels of residual lysosomal enzyme activity, down to a critical “threshold”, are sufficient to turn over substrate *in vivo* and prevent lysosomal storage disease and that small changes in residual activity below this threshold may lead to dramatic differences in the rate of substrate turn-over and the severity of the lysosomal storage disease (Conzelmann and Sandhoff, 1984). Several approved products for MPS have exploited this phenomenon to develop lysosomal-targeted enzyme replacement therapies (ERT) to augment residual lysosomal enzyme activity in patients, resulting in substrate reduction and improved quality of life (Poswar et al., 2017). These ERT trials have utilized the cell surface insulin-like growth factor 2 (IGF2) / cation-independent mannose-6-phospate receptor (CI-MPR) targeting pathway, where phosphorylated glycans present on secreted lysosomal enzymes bind avidly to the CI-MPR, resulting in their internalization into clathrin-coated vesicles, and delivery to lysosomes of patient cells (Kornfeld, 1992).

In the case of ERT development for Sanfilippo B, recombinant human NAGLU (rhNAGLU) over-expressed and secreted in Chinese hamster ovary (CHO) cells and in Sanfilippo B human fibroblasts is not mannose -6-phosphorylated (Zhao and Neufeld, 2000; Yogalingam et al., 2000), a property that may severely limit its uptake into key critical cellular targets of disease pathogenesis. Our group has therefore utilized a glycosylation independent lysosomal targeting (GILT) system, whereby IGF2 has been fused to rhNAGLU (rhNAGLU-IGF2; BMN 250) to mediate CI-MPR-mediated cellular uptake and delivery to lysosomes (LeBowitz et al., 2004, Maga et al., 2013; Kan et al., 2014). BMN 250 is a novel recombinant fusion protein currently being developed as an investigational ERT for Sanfilippo B. We have shown that intracerebroventricular (ICV) administration of BMN 250 in a knock-out mouse model and a naturally occurring dog model of Sanfilippo B can augment NAGLU activity and NAGLU protein levels in all cell types throughout the brain, normalize HS levels and reverse several aspects of well-entrenched secondary neuropathology associated with the disease (Kan et al., 2014; Aoyagi-Scharber et al., 2017; Ellinwood et al., 2018). While normalization of HS levels in the Sanfilippo B mouse and dog brains with ICV-administered BMN 250 is indicative of successful enzyme delivery to critical cellular targets of Sanfilippo B disease pathogenesis, the cellular uptake mechanism/-s by which this process occurs in different cell types of the brain has yet to be demonstrated.

Here we performed a series of competitive cellular uptake assays with BMN 250 in primary human Sanfilippo B patient fibroblasts and normal rodent-derived neurons, astrocytes and microglia. These studies distinguish two pathways by which BMN 250 in the extracellular space can be delivered to lysosomes of cells to augment residual NAGLU activity. We demonstrate that GILT technology confers predominantly CI-MPR-mediated cellular uptake and delivery of BMN 250 to lysosomes in neurons, astrocytes and fibroblasts, whereas receptor-independent cellular uptake in microglia contributes to substantial lysosomal delivery of both BMN 250 and the untagged rhNAGLU.

## METHODS

### Cell lines

Human MPSIIIB fibroblasts (GM02931) were obtained from the Coriell Institute for Medical Research (Camden, NJ). GM02931 were grown and passaged in MEM supplemented with 15% fetal bovine serum (FBS; 15% v/v) and nonessential amino acids (ThermoFisher Scientific, Waltham, M). Cortical neurons enriched from embryonic day 18 normal rat cerebrum were obtained from ScienCell Research Laboratories (Carlsbad, CA). Normal mouse-derived embryonic day 17 hippocampal neurons were obtained from Lonza Group Ltd. (Walkersville, MD). Astrocytes and microglia were enriched from postnatal day 2 normal rat total brain tissue were also obtained from ScienCell Research Laboratories (Carlsbad, CA). Neurons, astrocytes and microglia were cultured in poly-L-lysine-treated 96-well black tissue-culture-treated plates using defined medium, as recommended by the supplier.

### Direct “in-plate” NAGLU activity determination

NAGLU activity levels in cells plated in 96-well plates, as described above, was determined using the using the 4MU substrate, 4-Methylumbelliferyl-N-acetyl-α-D-glucosaminide. Briefly, cells in each well were washed extensively then lysed at room temperature for 15 minutes with 57.5 uL of mammalian protein extraction reagent per well (M-PER; ThermoFisher Scientific, Waltham, MA). Of this lysate, 10 uL was used to determine total protein levels using a BCA protein assay kit (ThermoFisher Scientific, Waltham, MA). The remaining lysate in each well was then incubated in the presence of 2 mM 4-Methylumbelliferyl-N-acetyl-α-D-glucosaminide in 0.2 M Acetate buffer pH 4.8 for 30 minutes at 37°C in a total reaction volume of 85 uL. The reactions were terminated by addition of 200 uL Glycine/NaOH buffer pH 10.7 to each well and plates were read at Ex360 Em460 with 455 cut off using a Spectramax i3 plate reader (Molecular Devices). A standard curve containing known amounts of 4-Methylumbelliferone (4-MU Standard; Sigma-Aldrich, St Louis, MO) was included in each assay to calculate NAGLU activity levels. In some instances where indicated, 4 uM IGF2 (Cell Sciences, Canton, MA) or 5 mM mannose-6-phophate (Man6P; Sigma-Aldrich, St Louis, MO) was added to uptake medium to competitively inhibit CI-MPR-mediated uptake of BMN 250.

### Enzyme uptake studies

Recombinant human NAGLU-IGF2 fusion and NAGLU proteins were expressed in Chinese hamster ovary cells and purified, as described elsewhere (US 9,376,480 B2; Kan et al., 2014). To determine how much BMN 250 is required to normalize NAGLU activity, endogenous NAGLU activity in each normal primary neuron, astrocyte or microglia cell line was compared with BMN 250-specific NAGLU activity following a 24 hour exposure to 6.25 nM BMN 250 or 6.25 nM rhNAGLU in the presence of 4 uM IGF2. Following BMN 250 uptake, cell lysates were prepared and assayed for NAGLU activity as described above. To determine the fold-increase above normal levels arising from BMN 250 uptake or rhNAGLU uptake, NAGLU activity following enzyme uptake was compared with the normal endogenous NAGLU activity level in each cell line.

### K_uptake_ determination

To determine the K_uptake_, the concentration of enzyme mediating half-maximal uptake, cells were incubated with varying concentrations of enzyme over a 24 hour period (dose-range 0.195 nM→100), at which time the enzyme activity present in cells was plotted into a Michaelis-Menten curve using GraphPad Prism software (La Jolla, CA).

### Immunofluorescence

BMN 250 uptake in primary cells was also monitored by imaging, where BMN 250 was directly conjugated with Alexa Fluor 488 (AF488) prior to uptake. BMN 250 was conjugated to AF488 using a 5 fold-molar excess of AF488 Sulfodichlorophenol Ester (Thermo Scientific A30052) and purified as described by the manufacturer, yielding approximately 3 fluorophores per molecule of BMN 250. NAGLU activity was tested before and after labeling to ensure that AF488 conjugation did not affect activity toward 4MU substrate (data not shown). Cells were incubated with 0.2 - 0.4 uM LysoTracker Red prior to fixation to permit identification of acidified lysosomes.

### Quantitative analysis of HS in MPSIIIB patient fibroblasts

Total heparan sulfate was quantified using a modification of the method previously described (Lawrence *et al.* 2012; Lawrence *et al.* 2008). Total HS data is expressed as average pmoles total HS per ug total protein from triplicate sample wells per condition +/- SD.

## RESULTS AND DISCUSSION

### BMN 250 uptake is predominantly CI-MPR-mediated in mouse hippocampal neurons

In normal mouse-derived hippocampal neurons, the majority of BMN250 uptake over 24 hours approximates Michaelis-Menten kinetics with a K_uptake_ of 5 nM (Figure 1A). AF-488-tagged BMN 250 co-localizes with LysoTracker Red^+^ acidified organelles in the cell body and along axonal projections of mouse neuronal hippocampal cells following cellular uptake, suggestive of successful delivery to lysosomes (Figure 1C, 1D and 1E). At the highest concentration tested (100 nM) NAGLU activity is increased by 13-fold, when compared with endogenous NAGLU activity detected in normal untreated mouse hippocampal neurons (Figure 1B). Furthermore, at this highest dose tested (100 nM) the majority of BMN 250 uptake in mouse hippocampal-derived neurons is inhibited in the presence of 4 uM IGF2, a competitive inhibitor of CI-MPR-mediated cellular uptake, suggesting that the majority of BMN 250 cellular uptake is CI-MPR-mediated (Figure 1B). CI-MPR-independent BMN 250 cellular uptake in mouse hippocampal neurons is sufficient to augment NAGLU activity by 2.5-fold, when compared with endogenous NAGLU activity detected in untreated normal mouse hippocampal neurons (Figure 1B). The untagged rhNAGLU is also taken up into mouse hippocampal neurons at the highest nM concentration tested (100 nM), which is sufficient to augment NAGLU activity by 2.1-fold above normal levels in untreated normal hippocampal neurons (Figure 1B). Glycan analysis of CHO-expressed NAGLU-IGF2 and untagged rhNAGLU molecules have previously demonstrated the absence of phosphorylated glycans (Kan et al., 2014). In support of this the addition of Man6P to the uptake medium does not inhibit uptake of rhNAGLU or BMN 250 (Figure 1B). These results suggest BMN 250 uptake is predominantly CI-MPR-mediated in mouse hippocampal-derived neurons, with receptor-independent uptake only becoming apparent at higher concentrations.

**Figure 1.**
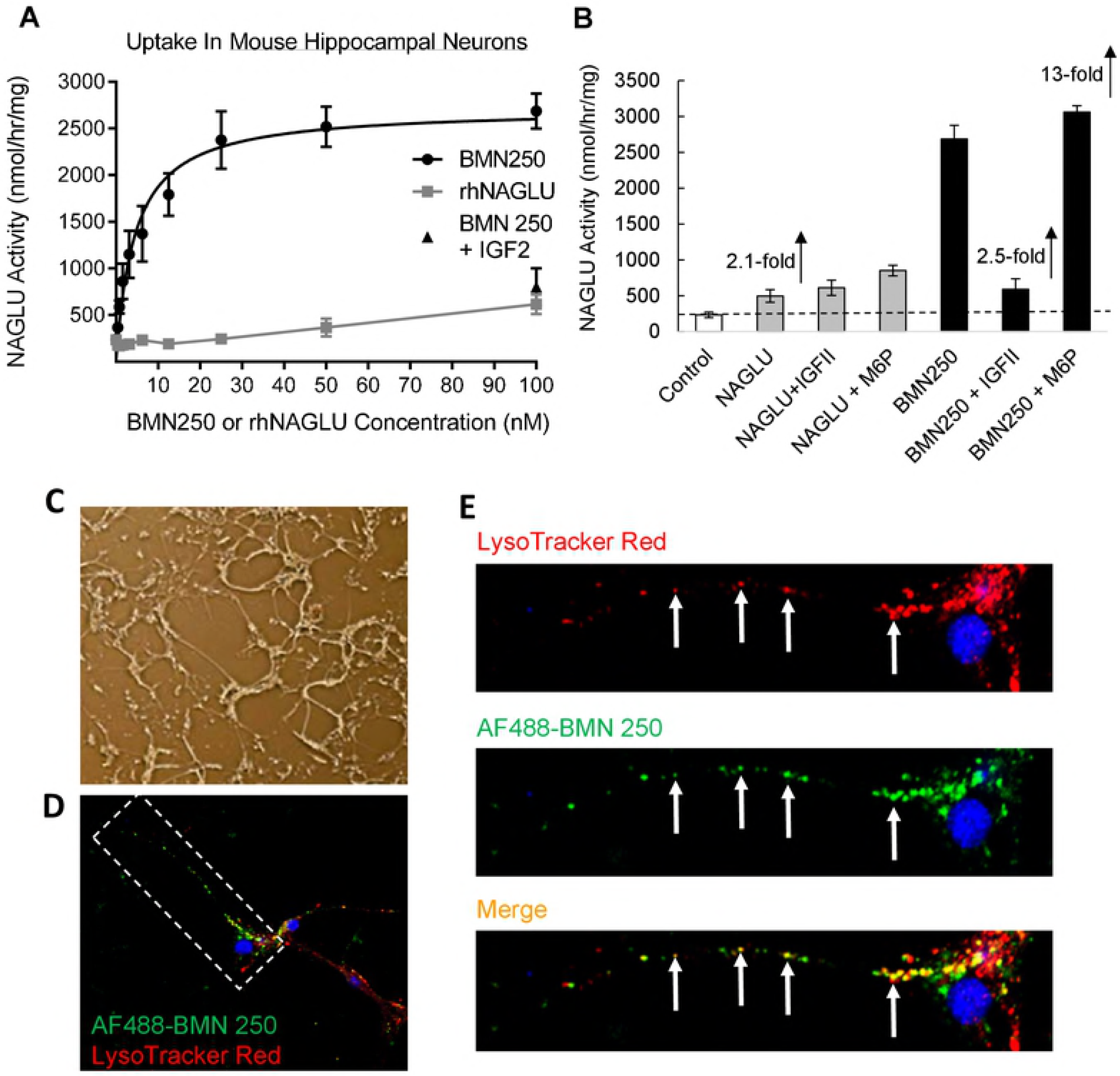
BMN 250 is delivered to lysosomes of mouse-derived hippocampal axonal projections. **A**, NAGLU activity detected in triplicate cultures of Sanfilippo B patient fibroblasts following uptake of BMN 250 and rhNAGLU over 24 hours. BMN 250 K_uptake_ = 4.5 nM; V_max_ = 2716 nmol/hr/mg. rhNAGLU K_uptake_ = 5.2 nM; V_max_=447 nmol/hr/mg **B**, NAGLU activity levels detected following uptake of rhNAGLU or BMN 250 in the absence or presence of 4 uM IGF2 or 5 mM M6P, as indicated. The level of endogenous NAGLU activity detected in normal mouse hippocampal neurons is indicated with a hashed line. **C**, Representative phase-contrast of cells described (B) following uptake with BMN 250; **D.** Merged confocal image of a single neuron incubated with AF488 BMN 250 and LysoTracker Red. **E.** Expanded images of the axonal projection highlighted in D. Arrows represent
examples of AF488 BMN 250 co-localization with LysoTracker Red^+^ organelles in the merged image.

### BMN 250 Uptake Into Rat Neurons and Astrocytes Is Predominantly CI-MPR-Mediated, Whereas Uptake In Microglia Occurs Through CI-MPR-Mediated and Receptor-Independent Mechanisms

BMN 250 cellular uptake was also evaluated in primary rat-derived neurons, astrocytes and microglia to permit comparison between three different cell types from the same species. NAGLU activity, standardized to total protein levels in each normal rat-derived cell line varies considerably, with comparatively higher levels of NAGLU activity being detected in normal rat-derived microglia (1120 ± 346 nmol/hr/mg), when compared with normal rat-derived cortical neurons (93 ± 23 nmol/hr/mg) and astrocytes (166 ± 12 nmol/hr/mg; Figure 2A). A low-dose exposure to 6.25 nM BMN 250 over 24 hours results in highly efficient cellular uptake of BMN 250 in all three primary rat cell lines, with 5-fold, 25-fold and 14fold increases in NAGLU activity above normal endogenous NAGLU activity levels detected in untreated rat neurons, astrocytes and microglia, respectively (Figure 2B, 2C, 2D). In contrast, NAGLU can only be detected in rat microglia following a 24 hour exposure to a low nM dose (6.25 nM) of the untagged rhNAGLU, with no NAGLU activity detected in rat derived neurons and astrocytes (Figure 2B, C, D, respectively). Furthermore, the NAGLU activity that was detected in rat microglia following exposure to a low nM dose of untagged rhNAGLU was increased by 2.5-fold over endogenous NAGLU activity detected in normal rat microglia (Figure 2D). These results suggest that of the three primary cell rat lines tested, BMN 250 can normalize NAGLU activity in all three cell lines at low nM concentrations, whereas untagged rhNAGLU can only normalize NAGLU activity in microglia.

**Figure 2.** Evaluation of BMN 250 and rhNAGLU uptake in normal rat neurons, astrocytes and microglia. **A.** Endogenous NAGLU activity levels present in normal rat-derived primary cortical neurons, astrocytes and microglia. To permit comparison between cell lines NAGLU activity levels present in cell lysates prepared from three separate cultures of each cell type was standardized to total protein levels and expressed as nmol/hr/mg total cell protein. Results are expressed as the mean (N=3) ± SD. **B, C, D**. NAGLU activity levels present in normal rat-derived primary cultures of cortical neurons (A), astrocytes (B) and microglia (C) following 24 hours of cellular uptake with 6.25 nM BMN 250 (black bars) or 6.25 nM rhNAGLU (grey bars). NAGLU activity is presented as the fold-increase above normal endogenous levels of NAGLU activity in each cell line.

To determine the enzyme uptake capacity (V_max_) for each rat-derived cell line a series of uptake curves were performed with increasing doses of BMN 250 or untagged rhNAGLU. As previously shown in ICV-ERT studies in Sanfilippo B mice (Aoyagi-Scharber et al., 2017), of the three rat-derived primary cell type cultures tested, untagged rhNAGLU only shows appreciable uptake in microglial cells, with low or non-detectable uptake detected in cortical neurons and astrocytes (Figure 3A). In contrast, BMN 250 uptake can be readily detected in all three cell types over a 24 hour period (Figure 3B), with K_uptake_ corresponding to 5 nM in cortical neurons, 3.4 nM in astrocytes and 2.6 nM in microglia (Figure 3B). These results are in agreement with our previous *in vivo* studies, with NAGLU protein being detected in neurons, astrocytes and microglia of the Sanfilppo B mouse brain following ICV administration of BMN 250 (Aoyagi-Scharber et al., 2017).

**Figure 3.** BMN 250 and rhNAGLU exhibit differential uptake capacities in normal rat-derived neurons, astrocytes and microglia. **A, B:** Representative curves of rhNAGLU uptake (A) or BMN 250 uptake (B) over 24 hours in triplicate cultures of normal rat-derived primary cultures of cortical neurons (squares), astrocytes (circles) and microglia (triangles). rhNAGLU k_uptake_ in microglia = 4.7 nM; rhNAGLU V_max_ in cortical neurons = 179 nmol/hr/mg, astrocytes = 371 nmol/hr/mg, microglia = 5469 nmol/hr/mg. BMN 250 K_uptake_ in cortical neurons = 5.05 nM, astrocytes = 3.4 nM, microglia = 2.6 nM. BMN 250 V_max_ in cortical neurons = 764 nmol/hr/mg, astrocytes = 5380 nmol/hr/mg, microglia = 9378 nmol/hr/mg.

BMN 250 uptake capacity in microglia (V_max_=12,336 nmol/hr/mg; Figure 3B) exceeds the uptake capacity reached in cortical neurons (V_max_=764 nmol/hr/mg; Figure 3B) and astrocytes (V_max_=5380 nmol/hr/mg; Figure 3B). The majority of BMN 250 uptake into rat cortical neurons and astrocytes is blocked with the addition of 4 uM IGF2 to the uptake medium, (Figure 4A, 4B), suggesting that BMN 250 uptake and delivery to lysosomes is predominantly CI-MPR-mediated. In contrast to rat neurons and rat astrocytes, rat microglia appear to possess a more efficient CI-MPR-independent uptake pathway, since the addition of 4 uM IGF2 to the uptake medium only partially inhibits the uptake of BMN 250 into rat microglia (Figure 4C). Receptor-independent uptake of BMN 250 in microglia plotted against enzyme concentration shows that the pattern of enzyme uptake does not reach saturation, further suggestive of receptor-independent uptake (Figure 5B). Likewise, cellular uptake of the untagged rhNAGLU in rat microglia is also observed (Figure 4C), which contrasts with rhNAGLU uptake in rat neurons and astrocytes (Figures 4A, 4B). These results are suggestive of both CI-MPR-mediated and receptor-independent mechanisms being utilized in the cellular uptake and lysosomal targeting of BMN 250 in rat-derived neurons, astrocytes and microglia, with the highest levels of receptor-independent uptake occurring in microglia.

**Figure 4.** BMN 250 exhibits CI-MPR-dependent and receptor-independent uptake in rat-derived neurons, astrocytes and microglia. **A, B and C:** Representative uptake curves of BMN 250 in the absence (black circles) or presence (grey squares) of 4 uM IGF2, or rhNAGLU (black triangles, hashed line) as indicated, in triplicate cultures of normal rat-derived primary cultures of cortical neurons (A; N=1 repeat) astrocytes (B; N=3 repeats) and microglia (C; N=3 repeats). BMN 250 K_uptake_ in rat cortical neurons = 6.0 nM, V_max_ = 1,040 nmol/hr/mg; BMN 250 K_uptake_ in astrocytes = 3.5 nM, V_max_ = 5,404 nmol/hr/mg; BMN 250 K_uptake_ in microglia = 2.6 nM, V_max_ = 12,336 nmol/hr/mg; BMN 250 K_uptake_ in microglia in the presence of 4 uM IGF2 = 19.7 nM, V_max_= 5028 nmol/hr/mg; rhNAGLU K_uptake_ in microglia = 4.1 nM; V_max_ = 5355 nmol/hr/mg.

**Figure 5.** Evaluation of CI-MPR-Dependent and Receptor-Independent BMN 250 Uptake In Rat-Derived Neurons, Astrocytes And Microglia. **A**, CI-MPR-dependent uptake of BMN 250 in triplicate cultures of rat microglia was calculated by subtracting endogenous NAGLU activity and the uptake achieved in the presence of 4 uM IGF2 from the total amount of uptake achieved with BMN250 in the absence of inhibitor. BMN 250 K_uptake_ in microglia = 2.2 nM, V_max_ = 9,009 nmol/hr/mg **B**, Receptor-Independent uptake of BMN 250 achieved in the presence of 4 uM IGF2 in microglia was calculated by subtracting endogenous NAGLU activity detected in untreated microglia. N = 3 repeats.

### BMN 250 Uptake In Sanfilippo B Patient Fibroblasts Is Predominantly CI-MPR-Mediated And Is Sufficient To Prevent HS Accumulation

Our results in primary neurons, astrocytes and microglia are suggestive of BMN 250 cellular uptake and delivery to lysosomes being mediated by both CI-MPR-mediated and receptor-independent mechanisms to varying degrees, depending on the enzyme concentration and the cell type. To further understand the therapeutic relevance of CI-MPR-mediated BMN 250 uptake we utilized Sanfilippo B patient fibroblasts that are known to accumulate HS (Prill et al., Submitted). Similar to the uptake pattern observed in neurons and astrocytes, BMN 250 uptake in Sanfilippo B patient fibroblasts over 24 hours approximates Michaelis-Menten kinetics with a K_uptake_ of 5.1 nM and is predominantly CI-MPR-mediated, since the majority of uptake can be inhibited with IGF2 (Figure 6). Receptor independent uptake is observed at the higher concentrations (Figure 6) and is similar to uptake of untagged rhNAGLU (Figure 6).

**Figure 6.** BMN 250 exhibits CI-MPR-dependent and receptor-independent uptake in Sanfilippo syndrome B patient fibroblasts. Representative uptake curves of BMN 250 in triplicate cultures of Sanfilippo B patient fibroblasts the absence (black circles) or presence (black squares) of 4 uM IGF2, or rhNAGLU (black triangles), as indicated. BMN 250 K_uptake_ = 5.1 nM; BMN 250 V_max_ = 5663 nmol/hr/mg. No NAGLU activity was detected in control untreated cells. N=3 repeats.

Incubation of Sanfilippo B patient fibroblasts with concentrations of BMN 250 at or below the 5.1 nM K_uptake_ (1.56, 3.125 or 6.25 nM BMN 250) for 2 hours, followed by 10 days of culture results in dose-dependent increases in NAGLU activity, corresponding to 8%, 16% and 21% of normal NAGLU activity levels detected in fibroblasts from a health control individual, respectively (Figure 7A). Uptake of BMN 250 coincides with dose-dependent reduction in the amount of HS accumulation (Figure 7A). Furthermore, at the highest concentration of BMN 250 tested (6.25 nM), HS levels are completely cleared, suggestive of successful delivery of BMN 250 to lysosomes of Sanfilippo B patient fibroblasts (Figure 7A). In support of this AF488-labeled BMN 250 co-localizes with LysoTracker Red^+^ organelles following cellular uptake in Sanfilippo B patient fibroblasts (Figure 7C). In contrast, untagged rhNAGLU does not exhibit cellular uptake in Sanfilippo B patient fibroblasts at the same concentrations tested (1.56 nM, 3.125 nM and 6.25 nM), which coincides with no reduction in HS storage (Figure 7B).

**Figure 7.** BMN 250 uptake in Sanfilippo syndrome B patient fibroblasts is sufficient to prevent HS accumulation. **A, B**, NAGLU activity present in Sanfilippo B patient fibroblasts incubated with doses of BMN 250 equivalent to or below the BMN 250 K_uptake_ (A; blue squares) or rhNAGLU (B; blue squares) for four hours on Day 2 of culture. Cells were washed extensively, then cultured for a further 8 days and analyzed for NAGLU activity using 4MU substrate. NAGLU activity detected is representative of three replicate cultures. Corresponding levels of HS detected at each enzyme concentration using SensiPro assay are indicated with black circles. NAGLU activity was augmented by 8%, 16% and 21% of normal NAGLU activity levels following incubation with 1.56 nM, 3.125, and 6.25 nM BMN 250, respectively, which coincides with near-to-complete clearance of HS. **C.** AF488-BMN 250 co-localizes with LysoTracker Red^+^ lysosomes following uptake in Sanfilippo B patient fibroblasts.

In conclusion, our results suggest that while untagged rhNAGLU exhibits efficient receptor-independent cellular uptake into microglia, negligible uptake of untagged rhNAGLU is observed in cultured neurons or astrocytes. BMN 250, an IGF2-tagged NAGLU molecule, overcomes this obstacle by permitting highly efficient CI-MPR-mediated cellular uptake and lysosomal delivery of NAGLU into neurons and astrocytes. Furthermore, only BMN 250 can mediate clearance of HS storage in primary cultures of Sanfilippo B mouse-derived neurons and astrocytes when exposed to very low nM concentrations (Prill et al., submitted), a phenomenon which may potentially occur at sites of the Sanfilippo B brain that are distal to the site of ICV-administered enzyme. If clearance of HS from neurons and astrocytes throughout the brain is critical for treatment of Sanfilippo B, then an efficient targeting mechanism such as that possessed by BMN 250 will be critical for successful treatment of this disease.

## Conflict of interest

The authors are employees of BioMarin Pharmaceutical, Inc. which is currently investigating BMN250 as a therapy for patients with MPSIIIB. Mika Aoyagi-Scharber, Terri M. Christianson and Jonathan H. LeBowitz, together with others, are inventors on a patent claiming compositions and methods for treating Sanfilippo type B, comprising therapeutic NAGLU-IGF2 fusion proteins (US 9,376,480 B2).

## Acknowledgement

We thank Gordon Vehar for his encouragement with initiating this exploratory project to support the BMN 250 program. Thanks also to Emma McCullagh for proof-reading the manuscript.

## Author Contributions

G.Y. carried out the study, directed cellular uptake studies and wrote the paper. A.L. performed cellular uptake assays. HP performed GAG analysis. M.J.L performed cloning, sequence verification, and generation of plasmid for all of the CHO transfections. B.Y. and J.H. performed protein purification. T.M.C. directed protein expression and production. M-A.S. was the BMN 250 project team leader, providing conceptual support. R.L. directed GAG analysis studies. B.E.C. contributed strategic input and supported the study. J.H.L. developed GILT tag technology, contributed strategic input, provided guidance with cellular uptake studies and directed the study.

## Abbreviations used

MPS: Mucopolysaccharidosis
NAGLU: alpha-N-acetylglucosaminidase
rhNAGLU: recombinant human NAGLU or tralesinidase alfa
IGF2: insulin-like growth factor 2
ERT: enzyme replacement therapy
CHO: Chinese hamster ovary
HS: heparan sulfate
HSPG: heparan sulfate proteoglycan
Man6P/M6P: mannose-6-phosphate
CI-MPR: cation-independent mannose 6-phosphate receptor
GILT: glycosylation independent lysosomal targeting

